# Ocular following responses of the marmoset monkey are dependent on post-saccadic delay, spatiotemporal frequency and saccade direction

**DOI:** 10.1101/2023.03.24.534057

**Authors:** Hoi Ming Ken Yip, Timothy John Allison-Walker, Shaun Liam Cloherty, Maureen Ann Hagan, Nicholas Seow Chiang Price

**Affiliations:** Department of Physiology and Neuroscience Program, Biomedicine Discovery Institute, Monash University, 26 Innovation Walk, Clayton, VIC, 3800, Australia; School of Engineering, RMIT University, Melbourne 3001, Australia

## Abstract

Ocular following is a short-latency, reflexive eye movement that tracks wide-field visual motion. It has been studied extensively in humans and macaques and is an appealing behaviour for studying sensory-motor transformations in the brain due to its rapidity and rigidity. We explored ocular following in the marmoset, an emerging model in neuroscience because their lissencephalic brain allows direct access to most cortical areas for imaging and electrophysiological recordings. In three experiments, we tested ocular following responses in three adult marmosets. First, we varied the gap between saccade end and stimulus motion onset (post-saccadic delay), from 10 to 300 ms. As in other species, tracking had shorter onset latencies and higher eye speeds with shorter post-saccadic delays. Second, using sine-wave grating stimuli we explored the dependence of eye speed on spatiotemporal frequency. The highest eye speed was evoked at ∼16 Hz and ∼0.16 cycles per degree (cpd), however, the highest gain was elicited at ∼1.6 Hz and ∼1.2 cpd. The highest eye speed for each spatial frequency was observed at a different temporal frequency, but this interdependence was not consistent with complete speed tuning of the ocular following response. Finally, we found the highest eye speeds when saccade and stimulus motion directions were congruent, although latencies were unaffected by direction congruence. Our results showed comparable ocular following in marmosets, humans and macaques, despite over an order of magnitude variation in body and eye size across species. This characterization will help future studies examining the neural basis of sensory-motor transformations.

**New & Noteworthy:** Previous ocular following studies focused on humans and macaques. We examined the properties of ocular following responses in marmosets in three experiments, in which post-saccadic delay, spatial-temporal frequency of stimuli and congruency of saccade and motion directions were manipulated. We have demonstrated short-latency ocular following in marmosets, and discuss the similarities across three species that vary markedly in eye and head size. Our findings will help future studies examining the neural mechanism of sensory-motor transformations.

## Introduction

Animals need to continually process sensory information and generate motor responses to interact with the environment. This sensory-motor transformation is still not well understood, even though in some circumstances it is highly stereotyped and can reliably occur within tens of milliseconds. The ocular following response is a smooth eye movement that almost immediately causes the eye to track the sudden movement of a large stimulus (Kawano & Miles, 1986; Miles et al., 1986; Miles & Kawano, 1986). It is useful for studying sensory-motor transformations because it is a stimulus-driven reflex that requires rapid sensory processing to extract visual motion information, which is then used to generate eye movements with a latency of just 60-80 ms. While there has been extensive research on ocular following in macaques (Adeyemo & Angelaki, 2005; Barthélemy et al., 2010; Ibbotson et al., 2007; Kawano & Miles, 1986; Miles et al., 1986; Miles & Kawano, 1986) and humans (Blum & Price, 2014; Gellman et al., 1990; Masson et al., 2000; Masson & Castet, 2002; Taki et al., 2009; Yang & Miles, 2003), it has not been examined in marmosets, a non-human primate animal model that has gained increasing interest in neuroscience research. The lissencephalic brain of marmosets makes them an attractive model because most brain areas are accessible on the cortical surface. In contrast, in macaques, many areas, like the middle temporal cortex, are buried in the sulci and difficult to reach. Our aim is to examine ocular following in marmosets, and characterise the stimulus properties that affect the latency and magnitude of the motor output. Parameterising the behavioural aspects of ocular following will help constrain future studies using neural recordings to explore sensory-motor transformations in the brain.

Ocular following is proposed to be a compensatory reflex that helps stabilise the gaze on the visual scene after a moving observer looks off to one side (Miles, 1998). It is closely related to the early component of the optokinetic response (Miles, 1998; Tabata et al., 2010), a stabilisation reflex evident in a wide range of species including those with less specialised cortical visual processing (e.g. rats and mice). Experimentally, a classical paradigm requires animals to first make a saccade from a peripheral position to a central target. After the saccade is completed, there is a delay period (i.e. post-saccadic delay) prior to the motion onset of a large-field visual stimulus in the background.

The eye tracking response is acutely stimulus-dependent. The onset latency is negatively correlated with stimulus speed and contrast, while the eye speed is positively correlated with these two properties (Miles et al., 1986). When tested with sine-wave gratings with a range of spatial frequencies and temporal frequencies, the highest eye speed was evoked at ∼0.4 cycles per degree (cpd) and ∼20 Hz in macaques (Miura et al., 2014). In humans, the optimal spatial frequency for eye speeds varied across individuals, but the optimal temporal frequency was consistently 16 Hz between observers (Gellman et al., 1990). In both humans and macaques, eye speeds have almost separable tunings for temporal and spatial frequencies of the stimuli, meaning that the preferred temporal frequency does not scale with the spatial frequency (Gellman et al., 1990; Miles et al., 1986; Miura et al., 2014). With increasing post-saccadic delay between saccade offset and motion onset of the stimulus, initial eye speeds (peak of eye speed at the first wave of eye acceleration) drop almost exponentially, while latency and later-phase eye speeds (averaged eye speeds at 100-140 ms) are less affected (Gellman et al., 1990; Kawano & Miles, 1986). The post-saccadic enhancement of eye speeds was suggested to be beneficial for suppressing post-saccadic ocular drifts (glissades).

Despite the short onset latencies, a multitude of cortical and subcortical areas are involved in generating a robust ocular following response. The pretectal nucleus of the optic tract (NOT) is a subcortical region that receives direct retinal projections and contains neurons predominantly tuned for tempero-nasal motion directions. Neurons in NOT fire before the ocular following response onset, and their firing rates are well predicted by sensory inputs (Inoue et al., 2000). Firing rates of individual neurons in the middle temporal area (MT) and the medial superior temporal area (MST) also ramp up before the ocular following responses start (Kawano et al., 1994) and depend on stimulus speed (Kawano et al., 1994) and post-saccadic delay (Ibbotson et al., 2007; Takemura & Kawano, 2006). Moreover, chemical lesions in MT and MST impair the magnitude of the eye-tracking response (Takemura et al., 2002, 2007), suggesting that these areas play a causal role in providing the sensory processing required for ocular following. It should be noted that the ocular tracking reflexes occur in species with less developed cortices, including rabbits (Collewijn, 1969) and mice (Tabata et al., 2010), and in these species, the role of the NOT is likely more important than that of MT/MST. Two subcortical regions, dorsolateral pontine nucleus (DLPN) and ventral paraflocculus (VPFL), are involved in the post-sensory processing of ocular following. DLPN is proposed to be an intermediate area that receives projection from MT/MST, computes further integration of information and project signals to VPFL. VPFL is suggested to be responsible for generating motor commands (Takemura & Kawano, 2002). Understanding how stimulus parameters affect neural activities, and eventually affect oculomotor response, would aid in elucidating the neural mechanism for sensory-to-motor transformations. Given that the essential sensory area MT is buried in the sulcus in macaques, marmosets are particularly suited for studying the neural processing of ocular following.

It is unclear whether short-latency ocular following occurs in marmosets, and if it does, how the gains and latencies compare with those in macaques and humans. Marmosets are known to engage in head tracking (Ngo et al., 2022), smooth pursuit eye movement (Mitchell et al., 2015), and eye tracking of optic flows (Knöll et al., 2018). It is thus reasonable to expect they exhibit the ocular following response, which is also a smooth eye tracking movement. However, their control of eye movement is poorer than macaques and humans, as reflected by lower gains and more frequent saccades in the smooth pursuit task (Mitchell et al., 2015). Therefore, it is important to characterise the differences between species.

In the current study, we tested three adult marmosets with a suite of stimuli that generate ocular following responses. We found reliable ocular following responses in all animals. Consistent with previous findings in humans and macaques, the latencies and speeds of ocular following responses in marmosets depend on stimulus properties. We found shorter latencies and higher eye speeds with shorter post-saccadic delays. The optimal spatiotemporal frequencies that evoked the highest eye speeds were similar to those found in humans and macaques, although the tuning for temporal frequency was more dependent on spatial frequency. We also explored the interaction of initial saccade direction and stimulus motion direction, which has not been systematically examined. It tells us how ocular following depends on the direction of the prior saccade, which is important for characterising the functional significance of ocular following. We found higher eye speeds when the two directions were congruent than incongruent. The results of this study will help future studies using advanced techniques such as multi-area electrophysiological recordings in the marmoset brain to disentangle the neural mechanisms underlying sensory-motor transformations (Zavitz & Price, 2019).

## Methods

### Animals

Three adult common marmosets (Monkeys Bu, Br, Ni, male, *Callithrix jacchus*) participated in this study. Monkey Ni was part of a previous study involving saccades and fixations. All procedures were approved by the Monash Animal Research Platform Animal Ethics Committee and followed the Australian Code of Practice for the Care and Use of Animals for Scienti?c Purposes.

### Surgery

We implanted a titanium head-post on each animal to stabilize its head during behavioural experiments. Surgery was performed under aseptic conditions. The animal was first injected with atropine (0.2 mg/kg, i.m.) and diazepam (2 mg/kg, i.m.). After 30 minutes, we induced anesthesia with alfaxalone (8 mg/kg, i.m.). The animal was placed in a stereotaxic frame and stabilised using earbars which had been covered in local anesthetic (2% xylocaine jelly). After intubation, the head was further stabilised with a palate bar and eye bars. Anesthesia was maintained by isoflurane (0.5-3%) in oxygen. Eyes were protected during surgery with liquid paraffin eye ointment. A midline incision was used to expose the skull and up to six titanium screws (diameter 1.5 mm, length 4 mm) were inserted 1-1.5 mm into the skull. The exposed skull was coated with a thin layer of dental adhesive (Supabond, Parkell). A head-post, which would stabilize the animal’s head during experiments, was then placed on the midline and transparent dental acrylic (Ortho-Jet; Lang Dental Mfg. Co.) applied to the base of the head-post and the screws to secure them to the skull. The margin was sealed with surgical adhesive (VetBond; 3M). Animals received oral antibiotics for 7 days (cefalexin monohydrate, 30 mg/kg) and analgesia for 5 days (meloxicam, 0.2 mg/kg). Monkey Bu and Br were also implanted with a titanium recording chamber in the same surgery, while Monkey Ni had a separate surgery to implant a chronic multi-electrode array in V1 of the right hemisphere.

### Stimulus and Procedure

Visual stimuli were generated in Matlab (MathWorks, Natick, MA) using Neurostim (https://klabhub.github.io/neurostim/) and the Psychophysics Toolbox extension (Brainard, 1997). Animals viewed the stimuli on a 24-inch Viewpixx/3D screen (refresh rate: 100 Hz; resolution: 1920 × 1080) with a fixed viewing distance of 56 cm.

We ran three experiments with identical procedures (Fig. 1). At the start of each trial, a stationary wide-field stimulus (a random dot pattern or sinusoidal grating with diameter 28 degrees), appeared on the screen with a peripheral annulus target (diameter 1 deg). The peripheral target could appear 5 degrees above, below, left, or right of the screen center. If the monkey’s eye stayed within 2 deg of the target for a random interval between 200 and 300 ms, the target would disappear and re-appear at the screen centre. Animals were required to make a saccade to this central target within 500 ms. If they successfully completed the saccade, the central target would disappear and after a post-saccadic delay period of up to 300 ms the wide-field stimulus would move for 300 ms. Stimulus motion was followed by an inter-trial interval of 1000 ms during which a juice reward was delivered. If the centering saccade was not completed within 500 ms, the trial was terminated with no reward.

**Fig. 1.**
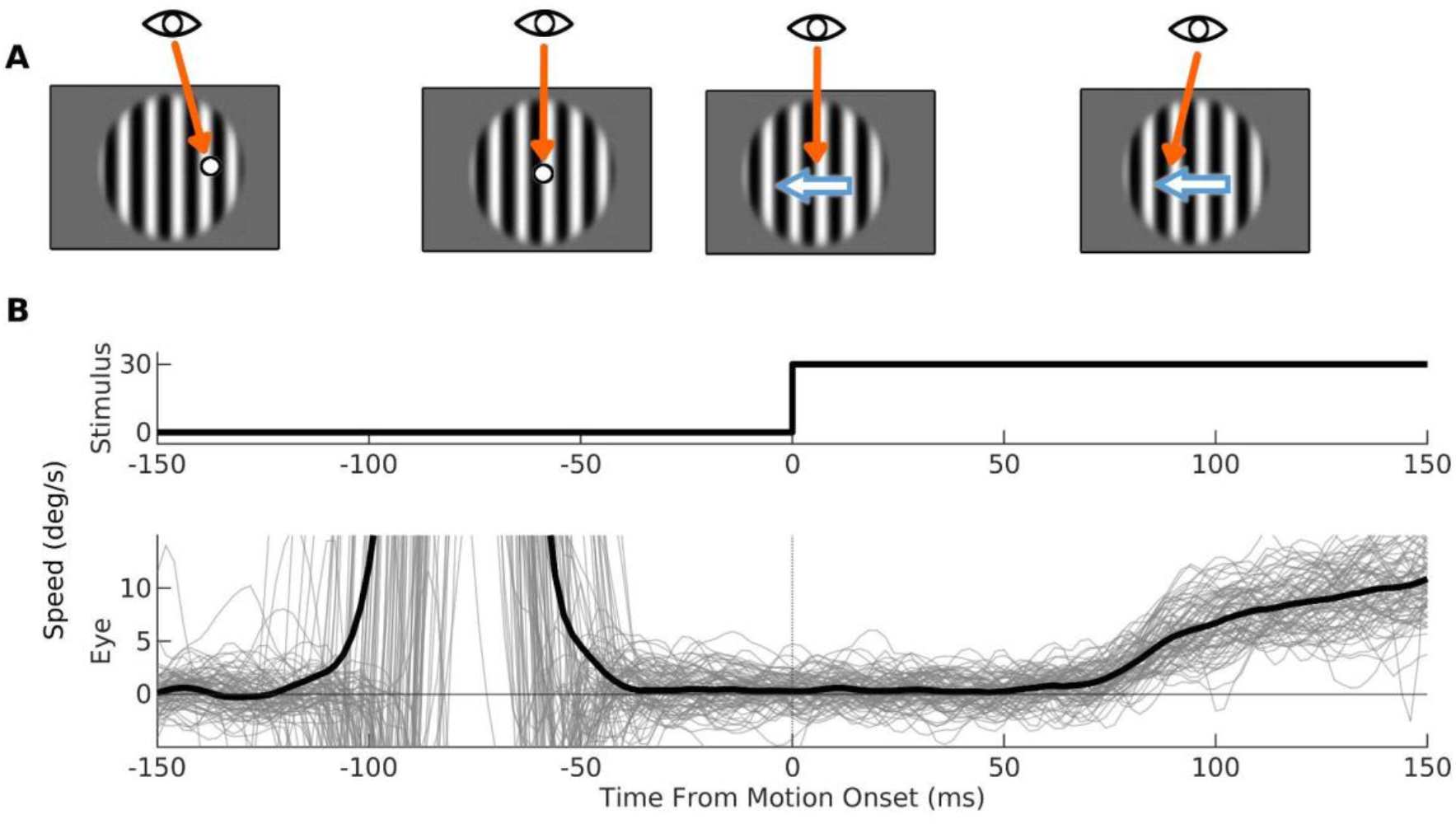
The ocular following paradigm. (A) Trials began with a large (28 deg) stationary stimulus (either a sinusoidal grating as shown or a random dot pattern) and a peripheral fixation target. Animals were required to fixate on the target for a randomized duration between 200 and 300 ms. Upon successful fixation, the target would disappear and reappear at the screen center. Animals were required to make a saccade to the new target within 500 ms. After the saccade, the stimulus would move following a post-saccadic delay (50 ms here). Eyes would track the stimulus motion soon after motion onset. (B) Stimulus and eye speed on 80 trials. For eye speed, the light grey lines represent data from individual trials and the black trace is the mean across trials.

In Experiment 1, we explored how post-saccadic delay affects tracking latency and speed. A random dot stimulus was used (2000 black dots with diameter 0.16 deg on a grey background, 100% coherence, speed 30 deg/s). We randomly varied the post-saccadic delay with 6 levels (10, 30, 50, 100, 200, 300 ms). Saccade and stimulus directions were either leftward or rightward and always congruent (e.g. the stimulus moved to the left following a leftward saccade).

In Experiment 2, we examined how spatiotemporal frequency affects eye speeds and gains. Stimuli were 100% contrast sinusoidal gratings with six temporal frequencies (1.56, 3.13, 6.13, 12.25, 18.75, 25 Hz) and seven spatial frequencies (0.04, 0.08, 0.16, 0.31, 0.62, 1.24, 2.48 cpd). These values were identical to a previous study in macaque (Miura et al., 2014), allowing us to make a direct comparison between species. The circular edge of the grating was masked with a cosine function to avoid sharp contrast changes at the edges. The post-saccadic delay was fixed at 50 ms. As in Experiment 1, saccade and stimulus directions were either leftward or rightward, and the two directions were always congruent.

In Experiment 3, we examined how the interaction between saccade and stimulus direction affects tracking latency and speed. The post-saccadic delay was fixed at 50 ms. We employed a random dot stimulus with the same parameters as in Experiment 1. Four saccade directions (upward, downward, leftward, rightward) were tested. For each saccade condition, we had two relative stimulus directions that were either congruent or incongruent with the saccade direction, e.g. leftward saccade was paired with leftward (congruent) or rightward (incongruent) stimulus motion.

### Eye velocity and latency

Eye movements were tracked with a video-based Eyelink 1000 system (SR Research). Horizontal and vertical eye positions were recorded in both eyes at 500 Hz sampling rates, although only data from one eye was used for analysis. At the start of each session, eye positions were calibrated by presenting small marmoset faces at different locations to encourage fixation. A Savitzky-Golay filter with order 3 and length 42 ms (21 data points) was used to smooth the position data and calculate eye velocity.

To estimate the onset latency at which the eyes started moving, we aligned eye velocity with stimulus motion onset in each trial. Then we averaged the eye velocity across trials within the same experimental conditions at each time point. Due to the noisiness of data at the single-trial level, estimating latency in individual trials gave us unreliable results. Therefore, we fitted a two-part piece-wise linear function (Equation 1) to the trial-averaged eye speed traces at 30 to 150 ms after motion onset with the least square method. The estimated value of *c* would therefore be our estimation of latency.

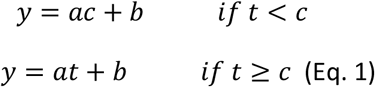

We employed a randomisation test to determine the significance of the relationship between post-saccadic delay and onset latency. First, we shuffled the condition labels for all trials and estimated the latency from the trial-averaged traces for each pseudo-condition. Then we found the slope of the simple linear regression model between latency and post-saccadic delay. By repeating this randomisation procedure 1000 times, we obtained a null distribution of the slope.

### 2D Gaussian Model

For each spatiotemporal frequency condition used in Experiment 2, we computed the trial-averaged eye speed in three time windows of interest (60 - 110 ms, 80 - 130 ms, 100 - 150 ms). We then fit a two dimensional Gaussian function (Equation 2) to the eye speeds using the same approach as a previous study in macaque (Miura et al., 2014).

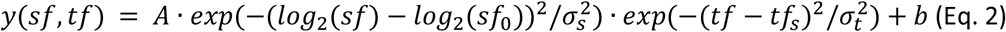

Where

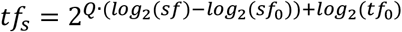

Parameters *A, b, sf*_0_, *tf*_0_, *σs, σt* are the gain, intercept, peak spatial frequency, peak temporal frequency, standard deviation of spatial frequency tuning and standard deviation of temporal frequency tuning, respectively. Parameter Q quantifies the degree of dependence on spatial frequency. If Q=1, the ocular following response is speed tuned; if Q=0, the ocular following response is temporal frequency tuned. Therefore, the lower and upper bound of Q were set at 0 and 1, respectively. To determine the significance of Q, we performed a F-test to compare the goodness of fit between a baseline model with Q=0 fixed and the full model with Q as a free parameter. A significant F-test result indicates that adding parameter Q improves the model fit.

## Results

### Experiment 1: Onset latencies and eye speeds depend on post-saccadic delay

In macaques and humans, the speed and onset latency of ocular following responses are strongly affected by post-saccadic delays, however, this has not been reported in other species. We first explored how post-saccadic delays affect the speed and latency of eye tracking in marmosets. At the start of each trial, a stationary, large-field random dot stimulus appeared in the background, with a small annulus target 5 deg left or right of the screen centre overlaid. Animals were required to fixate on the peripheral target for a randomized period of 200 to 300 ms, after which the target would disappear and reappear in the screen center. Animals were then required to make a saccade to the new target within 500 ms. Upon completion of a saccade, the background stimulus moved for 300 ms following post-saccadic delays of 10 to 300 ms (Fig 1). Eye speeds ramped up shortly after the motion onset (Fig 2A-C), although there was a large between-subject variation in gains. We found consistent post-saccadic enhancement in all animals: onset latencies and eye speeds were positively and negatively correlated, respectively, with the post-saccadic delay (Fig 2D-F).

**Fig. 2.**
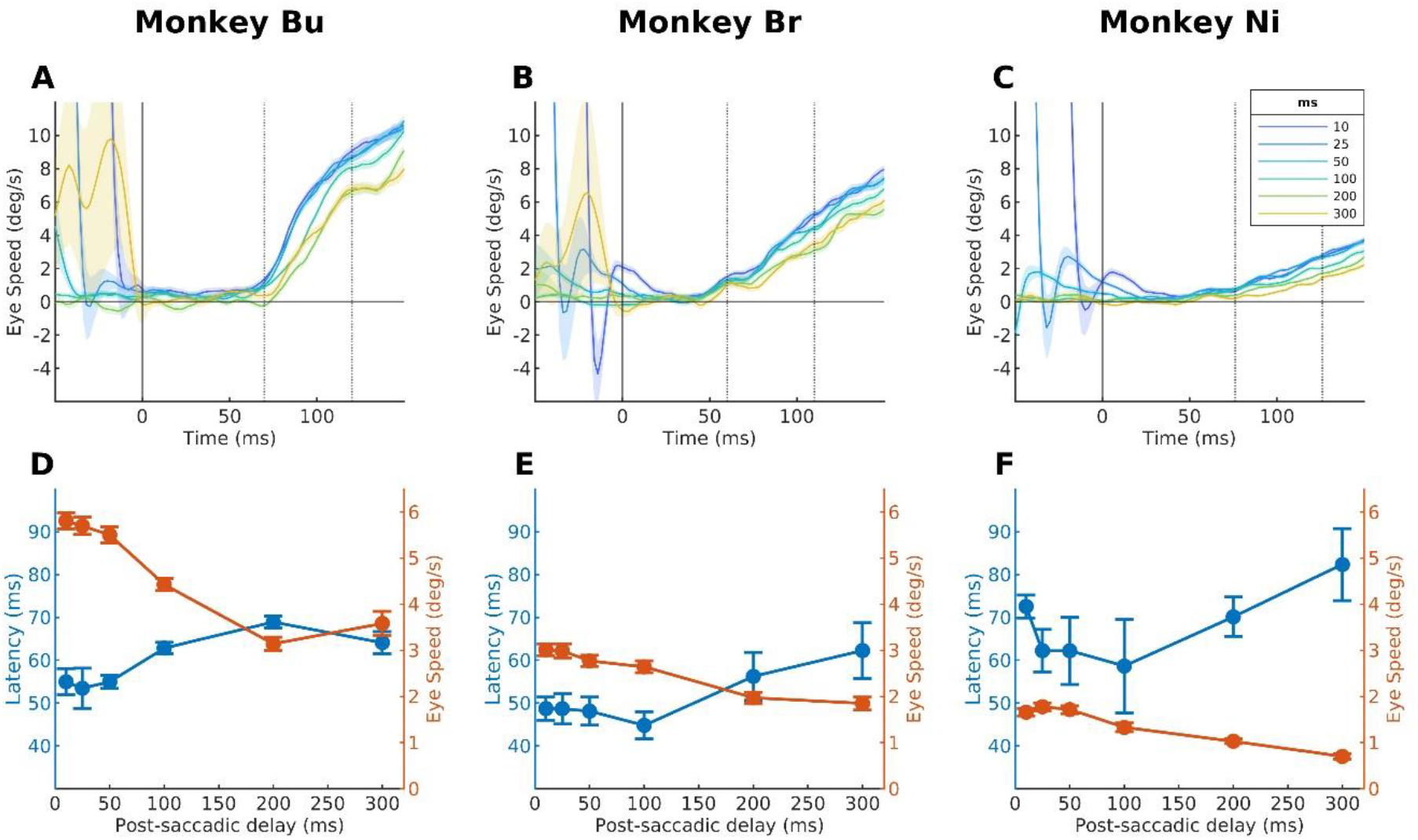
(A-C) Trial-averaged eye speeds over time for each animal. Each line shows the averaged eye speeds across trials for a single post-saccadic condition. Shaded region represents standard error of the mean across trials. Dashed lines indicate animal-specific time windows for averaging eye speeds. (D-F) Latency (blue) and eye speed (orange) for each post-saccadic delay. Error bars indicate standard deviation of the distribution of mean obtained with bootstrapping (n=1000).

We found significant positive correlations between onset latencies and post-saccadic delays in all animals (Fig 2D. Monkey Bu: slope = 0.045 ms/ms, p < 0.001; Fig 2E. Monkey Br: slope = 0.050 ms/ms, p < 0.001; Fig 2F. Monkey Ni: slope = 0.052 ms/ms, p = 0.013). Onset latencies were determined by fitting a piecewise linear function (Equation 1) to the trial-averaged eye speed traces for each condition. We employed a randomisation test to determine the significance of the relationship between post-saccadic delay and onset latency (see Methods). Comparison of the empirical slope and null distribution showed significant positive slopes in all animals.

Eye speeds were negatively correlated with post-saccadic delays. Mean horizontal eye speed was calculated in 50 ms time windows starting 10 ms after the median onset latency across conditions for each animal (Monkey Bu: 70 - 120 ms; Monkey Br: 60 - 110 ms; Monkey Ni: 76 - 126 ms). We then conducted a randomization test as described for latencies, which showed a significant slope in the relationship between post-saccadic latency and eye speed in all animals (Monkey Bu: slope = -9.13 deg/s^2^, p<0.001; Monkey Br: slope = -4.34 deg/s^2^, p<0.001; Monkey Ni: slope = -3.72 deg/s^2^, p<0.001). A limitation of calculating eye speed in a fixed time window is that the eye is moving slower at the start of the averaging window when the post-saccadic delay is longer. To address this, we also determined that the relationship between post-saccadic delay and eye speed had significant regression and slope when eye speed was calculated in a 50-ms window defined relative to the mean latency for each post-saccadic delay condition (Monkey Bu: slope = -2.89 deg/s^2^, p<0.001; Monkey Br: slope = -1.65 deg/s^2^, p=0.0284; Monkey Ni: slope = -2.46 deg/s^2^, p<0.001). To conclude, higher eye speeds and shorter latencies were found with shorter post-saccadic delays, which is consistent with findings in humans and macaques.

### Experiment 2: Eye speeds and gains depend on spatiotemporal frequencies

Previous studies in humans and macaques found a strong dependence of eye speed on spatiotemporal frequency. Therefore, we examined how the spatial and temporal frequency of a grating stimulus affects eye speed and gain. The procedure was identical to experiment 1 except we presented sine-wave gratings instead of random dot stimuli. We selected seven spatial (0.04 - 2.48 cpd) and six temporal frequencies (1.56 - 25 Hz), giving us 42 stimuli in total. The high-speed conditions (spatial frequency <= 0.31 cpd and temporal frequency >= 6.13 Hz) failed to trigger ocular following in Monkey Ni in most trials. With less than twenty valid trials in those conditions, it was hard to estimate eye speeds reliably. Therefore, Monkey Ni’s results were not reported here. In Monkey Bu and Br, the spatiotemporal profiles of eye speeds and gains were consistent across time (Fig 3).

**Fig. 3.**
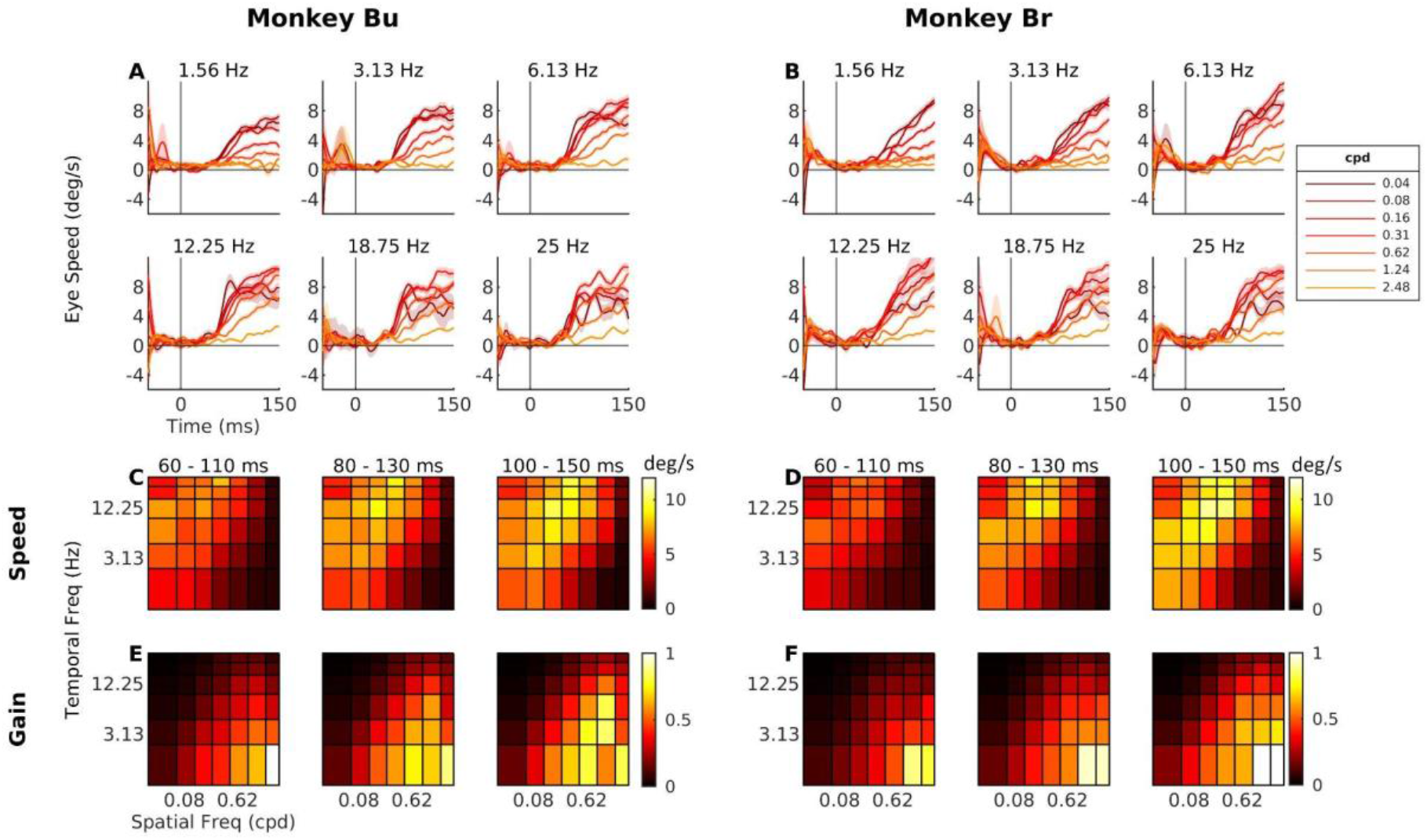
(A, B) Trial-averaged eye speeds over time for each animal separately plotted for each temporal frequency condition. Each line represents a different spatial frequency (cycles per degree, cpd). Darker color indicates lower spatial frequency. (C, D) Heat maps of eye speeds at three 50-ms time windows (60-110 ms, 80-130 ms, 100-150 ms). Each cell represents a spatiotemporal condition, and the color indicates eye speed. X- and Y-axes are in log scale. (E, F) Heat maps of gains at three 50-ms time windows. Gain is defined as eye speed divided by stimulus speed.

Since it was unclear how the spatiotemporal frequency tuning or the optimal frequency shift over time, we examined eye speeds in three time windows (60 - 110ms, 80 - 130 ms, 100 - 150 ms) in our analysis. The optimal spatial and temporal frequencies, which evoked the highest eye speeds, were consistently ∼0.16 cpd and ∼16 Hz across three time windows in both animals (Fig 3C,E). To examine whether eye speeds were tuned to speed or separately to spatial and temporal frequency, we fit a 2D Gaussian model to each time window in each animal and examined the parameter which quantifies the degree of dependence on spatial frequency (Q in the model, see *Equation 2*). A value of Q=0 means the spatial and temporal frequency tuning are separable, and a value of Q=1 indicates that the data are tuned to speed independent of spatial frequency. The estimated values of Q, in the order of early to late time windows, were 0.263, 0.246 and 0.215 in Monkey Bu and 0.225, 0.225 and 0.268 in Monkey Br respectively. The model had good fits in all time windows in both animals (R^2^ > 0.890). To examine the significance of Q, we conducted a F-test, which tests whether the full model fits the data better than a reduced model with fixed Q=0. All Q-values were significantly different from 0 under the F-test (p<.001), suggesting interdependence between tunings of spatial and temporal frequency.

Gain is defined as the ratio of eye speed and stimulus speed. The optimal spatial and temporal frequencies associated with the highest gains were consistent across time in both animals at ∼1.24 cpd and ∼1.56 Hz (Fig 3D,F). We did not fit the 2D Gaussian model to gains because the estimate of Q is hard to interpret when the optimal stimulus is close to the edge of the range of tested conditions. Gains were highest with low stimulus speeds (high spatial frequencies and low temporal frequencies), suggesting that it is challenging to choose a stimulus that produces oculomotor responses with both high gain and high absolute speed.

### Experiment 3: Eye speeds, but not latencies, depend on the congruence of saccade and stimulus direction

While the ocular following paradigm commonly involves a saccade preceding stimulus motion onset, it is not clear how the saccade and stimulus directions interact. Examining this interaction is important for understanding how ocular following depend on the prior saccade. We tested ocular following with four saccade directions (up, down, left, right), each with two motion directions that are congruent or incongruent with the saccade direction. For example, a leftward saccade direction could be paired with leftward (180°; congruent) or rightward (0°; incongruent) stimulus motion. We averaged the results from conditions with saccade directions in the horizontal and vertical axes after inverting the eye speeds in conditions with stimulus directions of 180 and 270° (Fig. 4A-C). Eye speeds were higher when saccade and stimulus directions were congruent than incongruent, for both vertical and horizontal dimensions. However, direction congruency had no effect on latencies.

**Fig. 4.**
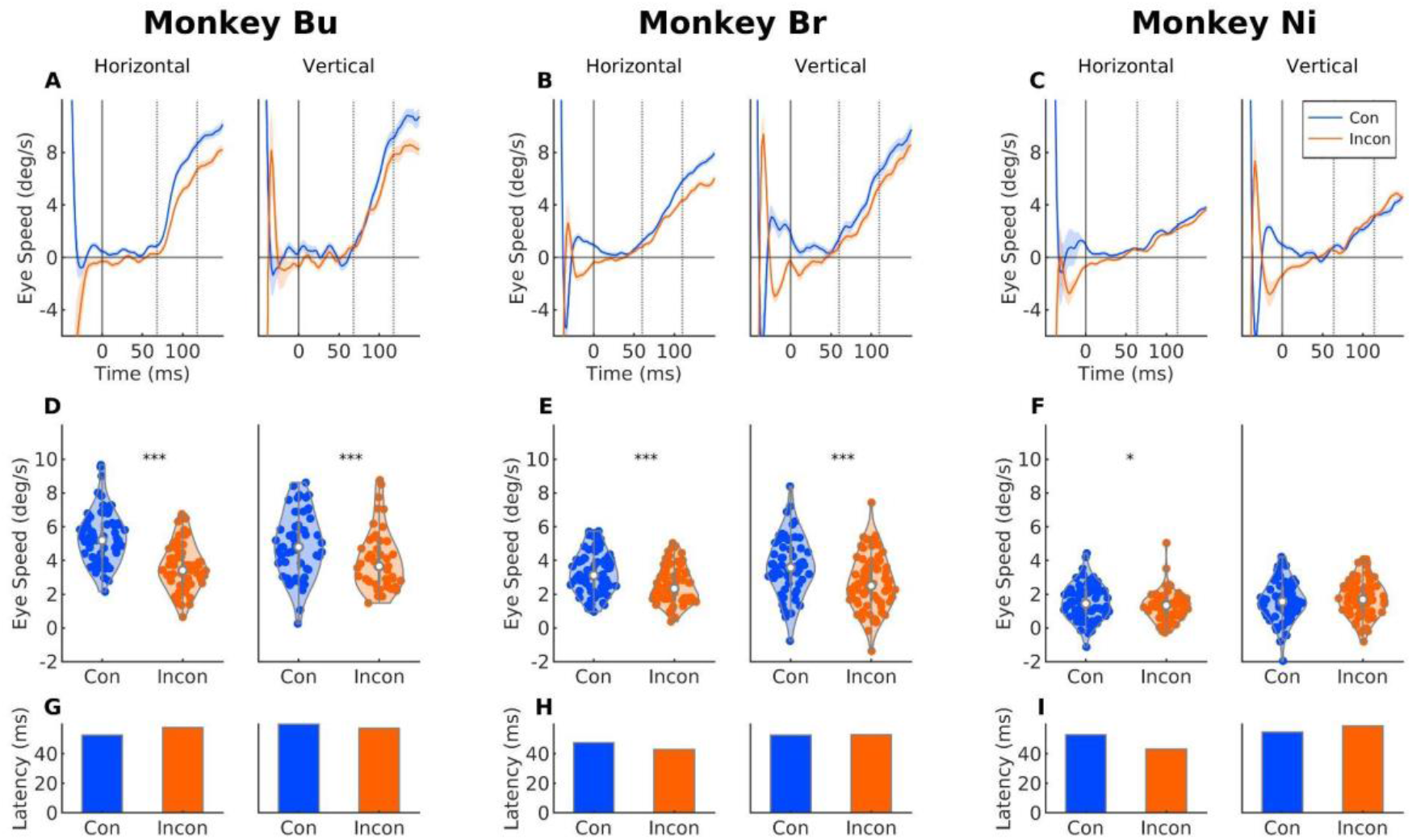
(A-C) Trial-averaged eye speeds for each animal for congruent (blue) and incongruent (orange) conditions, shown separately for horizontal and vertical dimensions. Dashed lines indicate time windows used to calculate eye speeds. (D-F) Comparison of eye speeds in congruent and incongruent conditions. Each dot represents the mean eye speed in a single trial. Asterisks indicate the significance level (*** p < 0.001; * p < 0.05) (G-I) Comparison of latencies in congruent and incongruent conditions. No conditions were associated with significant differences in latency.

Eye speeds generally depended on direction congruence. For Monkey Bu, mean eye speeds were significantly higher in congruent than in incongruent conditions in both horizontal (Fig 4D. t(62) = 6.08, p <0.0001) and vertical (Fig 4D. t(46) = 3.90, p <0.001) dimensions. A similar enhancement of congruency was also observed in Monkey Br (Fig 4E. horizontal: t (72) = 3.80, p<0.001; Fig 4E. vertical: t(62) = 3.92, p<0.001). For Monkey Ni, the congruency effect was significant in the horizontal dimension (Fig 4F. t(74) = 2.04, p=0.045) but not in the vertical dimension (Fig 4F. t(72) = - 0.698, p=0.487). As in Experiment 1, eye speeds were calculated as the mean amplitude in animal-specific 50 ms time windows (Monkey Bu: 68 - 118 ms; Monkey Br: 60 - 110ms; Monkey Ni: 64 - 114 ms).

There were no significant differences in latencies between congruent and incongruent conditions in both horizontal (Fig 4G. Monkey Bu: 5.196 ms, p =0.142; Fig 4H. Monkey Br: -4.558 ms, p = 0.136; Fig 4I. Monkey Ni: -9.581 ms, p = 0.226) and vertical dimensions (Fig 4G. Monkey Bu: - 2.908 ms, p = 0.174; Fig 4H. Monkey Br: 0.386 ms, p = 0.471; Fig 4I. Monkey Ni: 4.004 ms, p = 0.348). As in Experiment 1, we employed a randomization test to test significance. We shuffled the condition labels 1000 times, estimated latency of each pseudo condition, and calculated the latency difference between conditions. This formed a null distribution of latency difference, which we compared with the empirical values to assess significance.

## Discussion

In this study, we found three main results. First, the ocular following response in marmosets showed post-saccadic enhancement, reflected by shorter latencies and higher eye speeds with shorter post-saccadic delays. Second, eye speeds and gains were dependent on the spatiotemporal frequencies of the stimuli. Finally, congruent saccade and stimulus directions evoked higher eye speeds than the incongruent conditions, while direction congruence had no significant effect on latencies.

Our findings in Experiment 1 were qualitatively similar to previous reports in macaques and humans. First, the ocular following responses have ultra short latencies, although the latencies are even shorter in marmosets (∼50 ms) than macaques (∼60 ms) and human (∼80 ms). This is not surprising given that marmosets have considerably smaller eyes and brains, with the corresponding shorter axons suggesting faster responses are possible. Second, our three monkeys show markedly different gains for the random dot patterns with identical parameters. Likewise, previous macaque and human studies (Gellman et al., 1990; Miles et al., 1986), have also reported large inter-individual variations in gains for ocular following responses. Human studies of smooth pursuit eye movements have also found dramatic differences in gains across individuals (Bargary et al., 2017). Therefore, the gain in oculomotor system in primates, including marmosets, seems to exhibit large individual differences. Lastly, the dependence of eye speed on post-saccadic delay suggests that the ocular following response in marmosets may have a similar functional role to that hypothesised in humans and macaques. Eyes do not always stop at a designated target at the end of saccade and may instead drift forward or backward. This ocular drift, which is called a glissade, reduces the stability of the visual scene on the retina and reduces spatial acuity. By enhancing the gain of ocular following and shortening the latency until it begins in the period shortly after saccades, the visual system can suppress glissades (Ibbotson et al., 2007; Kawano & Miles, 1986).

The spatiotemporal frequencies that evoked the highest eye speeds in marmosets were similar to those previously reported in macaques and humans. In our study, gratings of ∼0.16 cpd and ∼16 Hz were optimal, compared with ∼0.4 cpd and ∼20 Hz in macaques (Miura et al., 2014). The lower optimal spatial frequency in marmosets than macaques is consistent with previous comparison of tunings in MT cells in the two primates (Lui et al., 2007). The higher prevalence of MT cells tuned to lower spatial frequencies in marmosets may be explained by the smaller eyes in marmosets, which have fewer retinal cells and hence lower spatial resolution. The optimal spatial frequency in humans varied a lot across individuals and thus no reliable estimates have been reported, while the optimal temporal frequency is consistently ∼16 Hz across observers (Gellman et al., 1990). It is not clear why the optimal temporal frequencies are similar across species. However, it should be noted that due to the 100 Hz refresh rate of our monitor, we did not present gratings with temporal frequencies higher than 25 Hz. Therefore, there is a possibility that the optimal temporal frequency in marmosets actually sits outside the range we tested.

Our findings suggested that the ocular following responses in marmosets are more speed-tuned than in macaques and humans. Tunings of temporal and spatial frequency are mostly separable in macaques, as reflected by an estimated value of 0 for the parameter Q in the 2D Gaussian model (Miura et al., 2014). There is no human data reported with 2D Gaussian fitting.

However, we could fit the model with the results reported from an experiment testing the ocular following response with gratings of 5 spatial frequencies and 5 temporal frequencies (Gellman et al., 1990). Using the data from two subjects, we obtained Q-values of 0.0261 and 0.144, which were not significantly different from 0 under the F-test. Our estimated Q-values (0.215 - 0.268) were higher, meaning our results contained more speed-tuning in responses than has been reported macaques and humans. Moreover, the 2D Gaussian model with Q as a free parameter had significantly better fits than the baseline model with Q-value fixed at zero, which suggests tunings of temporal and spatial frequencies are not completely separable. Previous studies in the tuning of MT cells in macaques revealed that a lot of neurons had at least intermediate tuning for temporal and spatial frequency (Perrone & Thiele, 2001; Priebe et al., 2003). However, most MT cells in marmosets showed separable selectivity for temporal and spatial frequency (Lui et al., 2007), which is contradictory to our findings in the oculomotor response. Future electrophysiological studies will be needed to clarify the relationship between stimulus representations in MT and ocular following behaviour.

The enhancement of eye speeds with congruent saccade and stimulus directions is consistent with findings on the effect of saccade-like stimulus, which is a high-speed stimulus that mimics the retinal input during saccade. By introducing a saccade-like stimulus before a test stimulus that actually triggers ocular-following response, It was found that eye speeds were higher when the saccade-like stimulus moves in the opposite direction of the test stimulus (Kawano & Miles, 1986).

Since a real saccade with the congruent direction as the stimulus creates retinal input similar to saccade-like stimulus in the opposite direction, our findings were consistent with this study. A possible explanation for this phenomenon is the involvement of attentive pursuit (Sheliga et al., 2005). When the saccade is in the same direction as stimulus motion, it may be assumed that the saccade serves to foveate a particular feature, which triggers the attentive pursuit system (Krauzlis, 2004). However, in our paradigm the saccade occurs over a stationary stimulus, which means attention can only take into effect within a short time window after saccade onset and before it ends.

A limitation of this study is that we did not test the full range of stimulus properties that have been examined in the ocular following studies in macaques. Instead, we replicated the major features of ocular following, including post-saccadic enhancement and dependence on spatiotemporal frequencies. Our results suggest that other properties of ocular following, for example dependence on eccentricity (Gellman et al., 1990; Miles et al., 1986), binocular disparity (Masson et al., 2001; Takemura et al., 2000; Yang & Miles, 2003) and pattern motion (Barthélemy et al., 2010; Masson et al., 2000; Masson & Castet, 2002), may also replicate in marmosets.

Our findings will help future studies in examining the neural mechanism of ocular following. Comparative analyses showed that the basic response properties of MT cells are similar in marmosets and macaques (Lui & Rosa, 2015). Areas MT and MST are buried within the superior temporal sulcus in macaques. In marmosets, these areas are fully accessible on the cortical surface. Using marmosets as an alternative animal model, therefore, opens opportunities for utilising contemporary technologies, including extracellular recording with multi-electrode arrays, two-photon calcium imaging of population activities and precise inactivation of cells with optogenetics etc. Furthermore, our results provide a quantitative basis for parameter selection. If maximising eye speed is important, we recommend using stimuli with spatial frequency ∼0.16 cpd, temporal frequency ∼16 Hz, and post-saccadic delay less than 50 ms.

In summary, our results showed that ocular following responses in marmosets depend on post-saccadic delay, spatiotemporal frequency of the stimulus, and the congruence between saccade and stimulus direction. These results can be utilized to design experiments that investigate sensory-to-motor mechanisms in the marmoset monkey.

## Acknowledgements

This project was funded by the Australian Research Council (DP2101002107 to NP; DE180100344 to MH; DP210103865 to SC) and by the National Health and Medical Research Council of Australia (APP1185442 to MH). KY was funded by the Monash Graduate Scholarship.

Data and analysis scripts can be found on https://github.com/nPrice-lab/OcuFolMarmoPaper.git

## Reference

Adeyemo, B., & Angelaki, D. E. (2005). Similar Kinematic Properties for Ocular Following and Smooth Pursuit Eye Movements. Journal of Neurophysiology, 93(3), 1710–1717. https://doi.org/10.1152/jn.01020.2004

Bargary, G., Bosten, J. M., Goodbourn, P. T., Lawrance-Owen, A. J., Hogg, R. E., & Mollon, J. D. (2017). Individual differences in human eye movements: An oculomotor signature? Vision Research, 141, 157–169. https://doi.org/10.1016/j.visres.2017.03.001

Barthélemy, F. V., Fleuriet, J., & Masson, G. S. (2010). Temporal Dynamics of 2D Motion Integration for Ocular Following in Macaque Monkeys. Journal of Neurophysiology, 103(3), 1275–1282. https://doi.org/10.1152/jn.01061.2009

Blum, J., & Price, N. S. C. (2014). Reflexive tracking eye movements and motion perception: One or two neural populations? Journal of Vision, 14(3), 23–23. https://doi.org/10.1167/14.3.23

Brainard, D. H. (1997). The Psychophysics Toolbox. Spatial Vision, 10, 433–436. https://doi.org/10.1163/156856897X00357

Collewijn, H. (1969). Optokinetic eye movements in the rabbit: Input-output relations. Vision Research, 9(1), 117–132. https://doi.org/10.1016/0042-6989(69)90035-2

Gellman, R. S., Carl, J. R., & Miles, F. A. (1990). Short latency ocular-following responses in man. Visual Neuroscience, 5(2), 107–122. https://doi.org/10.1017/S0952523800000158

Ibbotson, M., Price, N., Crowder, N., Ono, S., & Mustari, M. (2007). Enhanced Motion Sensitivity Follows Saccadic Suppression in the Superior Temporal Sulcus of the Macaque Cortex. Cerebral Cortex, 17(5), 1129–1138. https://doi.org/10.1093/cercor/bhl022

Inoue, Y., Takemura, A., Kawano, K., & Mustari, M. J. (2000). Role of the pretectal nucleus of the optic tract in short-latency ocular following responses in monkeys. Experimental Brain Research, 131(3), 269–281. https://doi.org/10.1007/s002219900310

Kawano, K., & Miles, F. A. (1986). Short-latency ocular following responses of monkey. II. Dependence on a prior saccadic eye movement. Journal of Neurophysiology, 56(5), 1355–1380. https://doi.org/10.1152/jn.1986.56.5.1355

Kawano, K., Shidara, M., Watanabe, Y., & Yamane, S. (1994). Neural activity in cortical area MST of alert monkey during ocular following responses. Journal of Neurophysiology, 71(6), 2305–2324. https://doi.org/10.1152/jn.1994.71.6.2305

Knöll, J., Pillow, J. W., & Huk, A. C. (2018). Lawful tracking of visual motion in humans, macaques, and marmosets in a naturalistic, continuous, and untrained behavioral context. Proceedings of the National Academy of Sciences, 115(44), E10486–E10494. https://doi.org/10.1073/pnas.1807192115

Krauzlis, R. J. (2004). Recasting the Smooth Pursuit Eye Movement System. Journal of Neurophysiology, 91(2), 591–603. https://doi.org/10.1152/jn.00801.2003

Lui, L. L., Bourne, J. A., & Rosa, M. G. P. (2007). Spatial and temporal frequency selectivity of neurons in the middle temporal visual area of new world monkeys (Callithrix jacchus). European Journal of Neuroscience, 25(6), 1780–1792. https://doi.org/10.1111/j.1460-9568.2007.05453.x

Lui, L. L., & Rosa, M. G. P. (2015). Structure and function of the middle temporal visual area (MT) in the marmoset: Comparisons with the macaque monkey. Neuroscience Research, 93, 62–71. https://doi.org/10.1016/j.neures.2014.09.012

Masson, G. S., Busettini, C., Yang, D.-S., & Miles, F. A. (2001). Short-latency ocular following in humans: Sensitivity to binocular disparity. Vision Research, 41(25), 3371–3387. https://doi.org/10.1016/S0042-6989(01)00029-3

Masson, G. S., & Castet, E. (2002). Parallel Motion Processing for the Initiation of Short-Latency Ocular Following in Humans. The Journal of Neuroscience, 22(12), 5149–5163. https://doi.org/10.1523/JNEUROSCI.22-12-05149.2002

Masson, G. S., Rybarczyk, Y., Castet, E., & Mestre, D. R. (2000). Temporal dynamics of motion integration for the initiation of tracking eye movements at ultra-short latencies. Visual Neuroscience, 17(5), 753–767. https://doi.org/10.1017/S0952523800175091

Miles, F. A. (1998). The neural processing of 3–D visual information: Evidence from eye movements. European Journal of Neuroscience, 10(3), 811–822. https://doi.org/10.1046/j.1460-9568.1998.00112.x

Miles, F. A., & Kawano, K. (1986). Short-latency ocular following responses of monkey. III. Plasticity. Journal of Neurophysiology, 56(5), 1381–1396. https://doi.org/10.1152/jn.1986.56.5.1381

Miles, F. A., Kawano, K., & Optican, L. M. (1986). Short-latency ocular following responses of monkey. I. Dependence on temporospatial properties of visual input. Journal of Neurophysiology, 56(5), 1321–1354. https://doi.org/10.1152/jn.1986.56.5.1321

Mitchell, J. F., Priebe, N. J., & Miller, C. T. (2015). Motion dependence of smooth pursuit eye movements in the marmoset. Journal of Neurophysiology, 113(10), 3954–3960. https://doi.org/10.1152/jn.00197.2015

Miura, K., Inaba, N., Aoki, Y., & Kawano, K. (2014). Difference in Visual Motion Representation between Cortical Areas MT and MST during Ocular Following Responses. Journal of Neuroscience, 34(6), 2160–2168. https://doi.org/10.1523/JNEUROSCI.3797-13.2014

Ngo, V., Gorman, J. C., De la Fuente, M. F., Souto, A., Schiel, N., & Miller, C. T. (2022). Active vision during prey capture in wild marmoset monkeys. Current Biology, 32(15), 3423–3428.e3. https://doi.org/10.1016/j.cub.2022.06.028

Perrone, J. A., & Thiele, A. (2001). Speed skills: Measuring the visual speed analyzing properties of primate MT neurons. Nature Neuroscience, 4(5), Article 5. https://doi.org/10.1038/87480

Priebe, N. J., Cassanello, C. R., & Lisberger, S. G. (2003). The Neural Representation of Speed in Macaque Area MT/V5. Journal of Neuroscience, 23(13), 5650–5661. https://doi.org/10.1523/JNEUROSCI.23-13-05650.2003

Sheliga, B. M., Chen, K. J., FitzGibbon, E. J., & Miles, F. A. (2005). Initial ocular following in humans: A response to first-order motion energy. Vision Research, 45(25), 3307–3321. https://doi.org/10.1016/j.visres.2005.03.011

Tabata, H., Shimizu, N., Wada, Y., Miura, K., & Kawano, K. (2010). Initiation of the optokinetic response (OKR) in mice. Journal of Vision, 10(1), 13. https://doi.org/10.1167/10.1.13

Takemura, A., Inoue, Y., & Kawano, K. (2000). The effect of disparity on the very earliest ocular following responses and the initial neuronal activity in monkey cortical area MST. Neuroscience Research, 38(1), 93–102. https://doi.org/10.1016/S0168-0102(00)00149-8

Takemura, A., Inoue, Y., & Kawano, K. (2002). Visually Driven Eye Movements Elicited at Ultra-short Latency Are Severely Impaired by MST Lesions. Annals of the New York Academy of Sciences, 956(1), 456–459. https://doi.org/10.1111/j.1749-6632.2002.tb02854.x

Takemura, A., & Kawano, K. (2002). Sensory-to-motor processing of the ocular-following response. Neuroscience Research, 43(3), 201–206. https://doi.org/10.1016/S0168-0102(02)00044-5

Takemura, A., & Kawano, K. (2006). Neuronal responses in MST reflect the post-saccadic enhancement of short-latency ocular following responses. Experimental Brain Research, 173(1), 174–179. https://doi.org/10.1007/s00221-006-0460-4

Takemura, A., Murata, Y., Kawano, K., & Miles, F. A. (2007). Deficits in Short-Latency Tracking Eye Movements after Chemical Lesions in Monkey Cortical Areas MT and MST. Journal of Neuroscience, 27(3), 529–541. https://doi.org/10.1523/JNEUROSCI.3455-06.2007

Taki, M., Miura, K., Tabata, H., Hisa, Y., & Kawano, K. (2009). The effects of prolonged viewing of motion on short-latency ocular following responses. Experimental Brain Research, 195(2), 195–205. https://doi.org/10.1007/s00221-009-1768-7

Yang, D.-S., & Miles, F. A. (2003). Short-latency ocular following in humans is dependent on absolute (rather than relative) binocular disparity. Vision Research, 43(12), 1387–1396. https://doi.org/10.1016/S0042-6989(03)00146-9

Zavitz, E., & Price, N. S. C. (2019). Understanding Sensory Information Processing Through Simultaneous Multi-area Population Recordings. Frontiers in Neural Circuits, 12. https://www.frontiersin.org/article/10.3389/fncir.2018.00115

